# Development of appropriate sensorimotor recalibration for a de novo skill

**DOI:** 10.64898/2026.11.02.763836

**Authors:** Tianhe Wang, Yuling Li, Jordan A. Taylor, Richard B. Ivry

**Affiliations:** Princeton Neuroscience Institute, Princeton, New Jersey; Department of Psychology and Neuroscience, University of California, Berkeley, California; Department of Molecular and Cell Biology, University of California, Berkeley, California; Department of Psychology, Princeton University, Princeton, New Jersey

**Author notes:** Corresponding author: Tianhe Wang.

## Abstract

Implicit adaptation operates automatically to ensure that well-learned movements remain exquisitely calibrated. A long-standing puzzle is how this process is applied when learning a new skill: Prior work has highlighted that implicit recalibration is inflexible, even when its operation results in maladaptive behavior given the sensorimotor mapping required for the new skill. Here, we revisited this problem, training participants for up to a month on a range of tasks with novel mappings, including mirror reversals and large rotations. Contrary to previous findings, implicit recalibration was flexibly reconfigured after prolonged training in a manner appropriate for each novel mapping. Training that emphasized feedforward planning rather than feedback control was sufficient to induce this reconfiguration. Learning involved reconfiguration of the sensory predictions and error correction policy for the new mapping, and participants were able to switch between mappings with minimal interference. These results demonstrate that implicit recalibration is reconfigured during skill acquisition, enabling flexible, context-specific adaptation.

## Introduction

Humans and animals continuously adjust their motor output based on errors. One important process that supports this ability is sensorimotor recalibration, which, in an implicit, automatic manner, uses error information to keep the sensorimotor map exquisitely tuned. While this implicit recalibration system has been well characterized in simple reaching behavior^1–5^, it remains unclear how this system functions in motor skills where the relationship between error and the required response to that error is based on a different sensorimotor map^6–11^. For example, when the heading of a sailboat drifts slightly to the left of the desired direction, the tiller is pushed to the left, a correction that is counterintuitive. Some classic findings imply that the implicit system might be capable of accommodating novel mappings. For example, Stratton (1896) describes becoming facile in moving about the world after wearing inverting prisms for eight days^12^ and the amnesic patient H.M. showed clear day-by-day improvement on mirror-tracing despite forming no conscious memory of the practice^13^.

However, studies using rigorous probes of implicit recalibration have indicated that implicit recalibration is surprisingly rigid, showing little ability to adjust to novel sensorimotor mappings^6,7,14,15^. Much of this work comes from studies using visuomotor adaptation tasks as a model paradigm. When the perturbation involves a rotation or translation, adaptation entails an adjustment to the sensorimotor map that directly reflects hand movement, or what we will refer to as a default mapping; for example, adapting to a counterclockwise (CCW) rotation requires an adjustment of the movement heading in the clockwise (CW) direction (Fig 1a). In contrast, if the sensorimotor map is mirror-reversed (as with our tiller example), an adaptive correction to the same CCW rotation would require an adjustment of movement that is also in the CCW direction (Fig 1b). However, even after one week of training, implicit sensorimotor recalibration remained grounded in the default mapping^6^, even though participants could easily select the proper heading direction to reach to a visual target (i.e., the mirrored location). That is, the automatic response to an error signal was maladaptive, producing trial-by-trial corrections that were in the wrong direction for the mirror-reversed environment (Fig 1c). This maladaptive behavior has been replicated across several studies^7,8,11,16,17^, and to date no evidence has shown proper implicit adaptation under mirror reversal.

**Figure 1.**
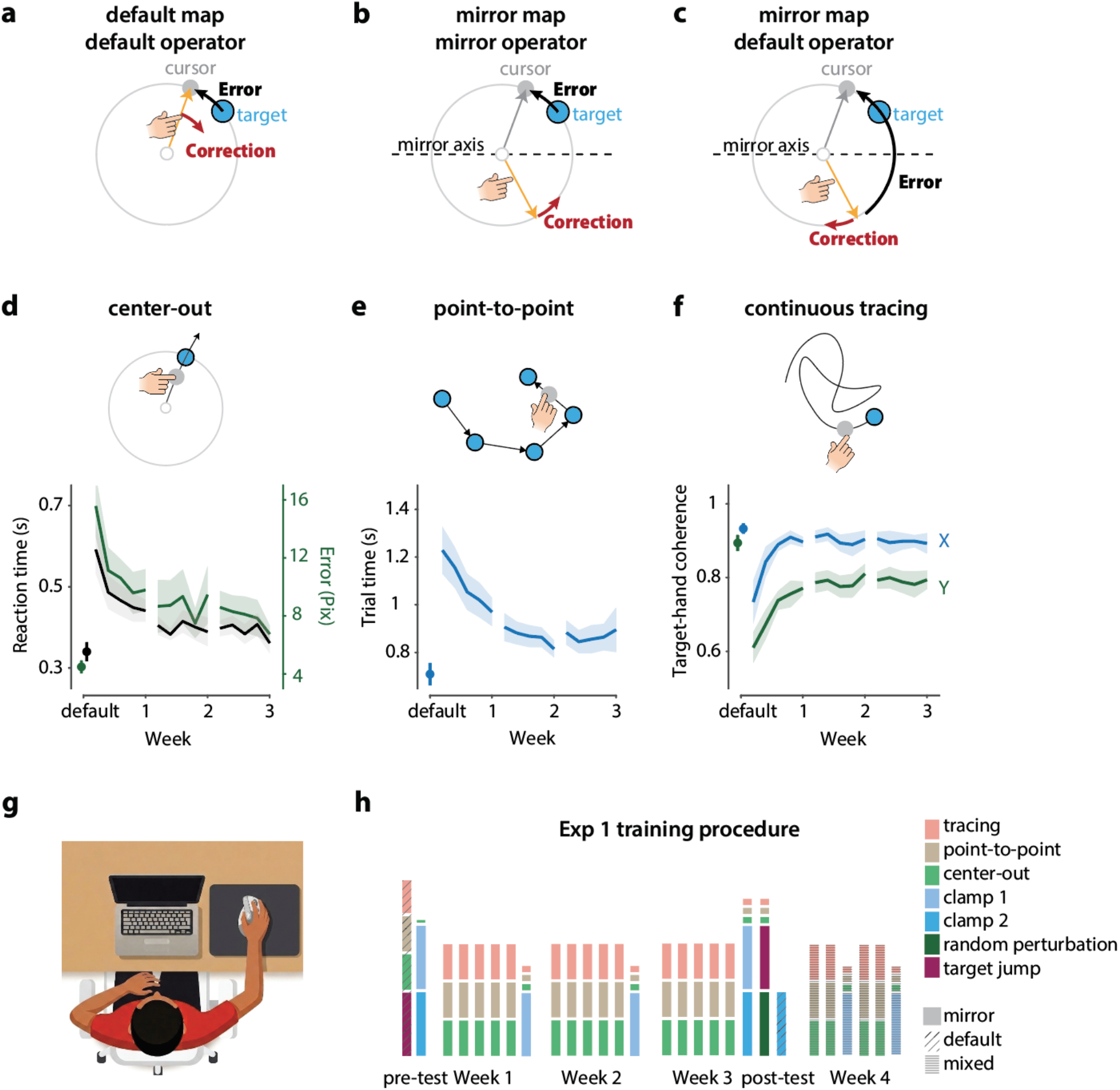
Learning during mirror-reversal training. (a–c) Illustration of error-correction direction in implicit recalibration for different visuomotor mappings. (a) In the default map, the default operator calculates error as the difference between the feedback cursor and the aimed hand position (here, the target). The correction is applied in the opposite direction of the error. (b) When the operator is adapted to the mirror map, the error is calculated as the difference between the feedback cursor and the predicted outcome of the mirror operation (target location). The correction is also mirror-reversed and now appropriate for the novel context. (c) When the default operator is applied to the mirror map, the error is also computed as the difference between the feedback cursor and the aimed hand position (mirror location of target), making the correction maladaptive. (d–f) Learning curves for the three training tasks. Dots preceding the curves indicate baseline performance under the default map. (d) Center-out reaching task: Participants sliced through a target appearing at a random location on an invisible circle. (e) Point-to-point reaching task: Participants reached to a series of targets with a fixed distance between consecutive targets. The next target appeared only after the previous one was reached. (f) Continuous tracing task: Participants traced a continuously moving target for one minute, controlling the position of a feedback circle. (g) Experimental setup. The laptop was positioned at the midline of the body, roughly 50 cm from the body. The mouse pad was positioned to the right of the laptop. With their upper limb resting on the table, the participant held a vertical mouse with a pinch grip. (h) Training procedure in Experiment 1. Each bar represents one day, with the exception of the pre-test assessments which were all performed on day 1. The post-test assessments after week 3 were performed across three consecutive days. The stacked segments show the tasks performed in that session, ordered from top to bottom by the order in which they were performed, with the stack roughly showing the training duration of each task. The training procedure diverged slightly across individuals (see Supplementary Methods). The two clamp types (scenarios 1 and 2) are defined in Fig 2.

We note three issues that qualify the inferences that can be drawn from prior studies on this problem. First, the training duration may have been insufficient for the implicit system to fully learn the new mapping. Second, previous studies have generally relied on center-out reaching tasks, a paradigm in which the workspace is limited^11^. Third, while these tasks have focused on feedforward control (e.g., trial-to-trial recalibration of the sensorimotor map), effective recalibration within a new sensorimotor map may require training in online feedback control, enabling the implicit system to learn how to perform immediate error correction based on the sensory feedback^18–21^.

Here we re-examine whether the implicit system can be properly reconfigured for a novel mapping, using a protocol designed to address these limitations. We recruited participants for a month-long study, training them to reach to visual targets in a mirror-reversed environment. We evaluated the operation of implicit recalibration by probing their response to an additional rotational perturbation that was imposed on top of the mirror map. To provide a rich training environment for learning the mirror-reversed mapping, the participants completed three tasks each day: center-out reaching, point-to-point reaching, and continuous tracing. With this set of tasks, the movements were produced over a large workspace and would be expected to engage both feedforward and feedback control processes, allowing us to evaluate the role of online control in reconfiguring the implicit system. Through this design, we sought to provide a strong test of the purported inflexibility of implicit recalibration: Will it continue to operate on a default sensorimotor map, and thus, yield maladaptive corrections? Or will it come to operate on a novel sensorimotor map, and thus be adaptive in support of a newly acquired skill?

## Results

### Mirror-reversal training

To examine if implicit recalibration can operate in a manner that is optimal for a novel sensorimotor map, in Exp 1, we trained participants on a variety of mirror-reversal tasks in which we inverted the relationship between movement and the visual feedback of that movement across the x-axis. In this manner, visual feedback of the vertical component of the movement was flipped (e.g., downward hand displacement resulted in upward cursor displacement). Feedback of the horizontal component of the movement was veridical (e.g., rightward hand displacement resulted in rightward cursor displacement).

Training was conducted over the course of one month (Fig 1h), well beyond the duration of previous laboratory studies. We used a diverse training schedule in which participants practiced three tasks (Fig 1d–f) each day, designed to promote both feedforward and feedback control (see Methods for details). To encourage precise, feedforward planning, the center-out reaching task required fast, ballistic movements, attempting to intersect a target with minimal opportunity for online corrections. To require demanding online control, the continuous tracing task required participants to continuously adjust their movements to follow an unpredictably moving dot^10,11^. The point-to-point reaching task required a series of discrete movements to points distributed across the workspace. This task was selected as an intermediate condition, requiring both feedforward planning for the initial movement heading and online corrections to reach each target. Targets and trajectories across all three tasks were designed to provide experience across a large portion of the available workspace.

Participants completed a 30-min training block each day over a 3-week training period (Fig 1g–h), with hybrid in-lab and at-home sessions, both using a MacBook. They gradually improved on all three tasks, demonstrating successful learning of the mirror-reversed mapping (Fig 1d–f). Change in performance was measured by fitting a linear mixed-effects model with log-transformed training day as the fixed effect. On center-out reaching, both absolute error (β = −2.3°, z = −5.3, p < 0.001) and reaction time (RT; β = −65 ms, z = −4.0, p < 0.001) decreased across training; point-to-point movement time shortened (β = −148 ms, z = −4.0, p < 0.001); and target–hand coherence improved along both the non-perturbed x-axis (β = 0.056, z = 2.5, p = 0.014) and the mirror-reversed y-axis (β = 0.065, z = 5.5, p < 0.001). However, even at the end of training, performance with the mirror map remained significantly worse than pre-training performance with the default map (all ts > 2.9, ps < 0.02, ηp² > 0.43, BF₁₀ > 4.0), a map acquired from a lifetime of experience. It remains an open question whether, and when, performance on the mirror map would reach a level comparable to that observed with the default map.

### Implicit recalibration is reconfigured after prolonged training of mirror reversal

Our primary focus is on the participants’ response to a feedback perturbation under a mirror-reversed mapping. In most motor tasks, feedforward error correction can be carried out by multiple processes including explicit, strategic aiming and implicit sensorimotor recalibration. To measure the operation of implicit recalibration, we used the clamped visual feedback technique (see Methods)^5,22^. Here the cursor follows a fixed angular trajectory relative to the target, independent of the participant’s hand direction. When performed under a default sensorimotor mapping, the heading direction gradually shifts in the direction opposite to the cursor even when participants are instructed to ignore it and believe they are reaching straight to the target (Fig 1a). As such, participants show signatures of implicit recalibration in this task even though the error remains invariant given the lack of contingency between the hand position and cursor.

In a mirror-reversed mapping, the same response following the default mapping could sometimes be maladaptive (Fig 1c), increasing the error on the following trial. To be adaptive in the mirrored context, two core computations would need to be modified (Fig 2a). First, the system would need to generate accurate sensory predictions of an action under the new mapping, predictions that can be compared with the feedback to compute a sensory prediction error (SPE)^4,23^. Under a mirror reversal, the system would learn to predict that the cursor moves in the mirror-reversed direction of the hand movement; thus, the error is computed relative to this mirror-reversed location, namely the target location. Second, the system would need to learn the appropriate error correction policy that maps sensory errors to motor commands^24,25^: in contrast to the policy under the default mapping where the proper correction is opposite the direction of the error, a policy under a mirror-reversed mapping would require that the correction be in the same direction as the error. We refer to the pairing of a sensory prediction and an error-correction policy as an operator.

**Figure 2.**
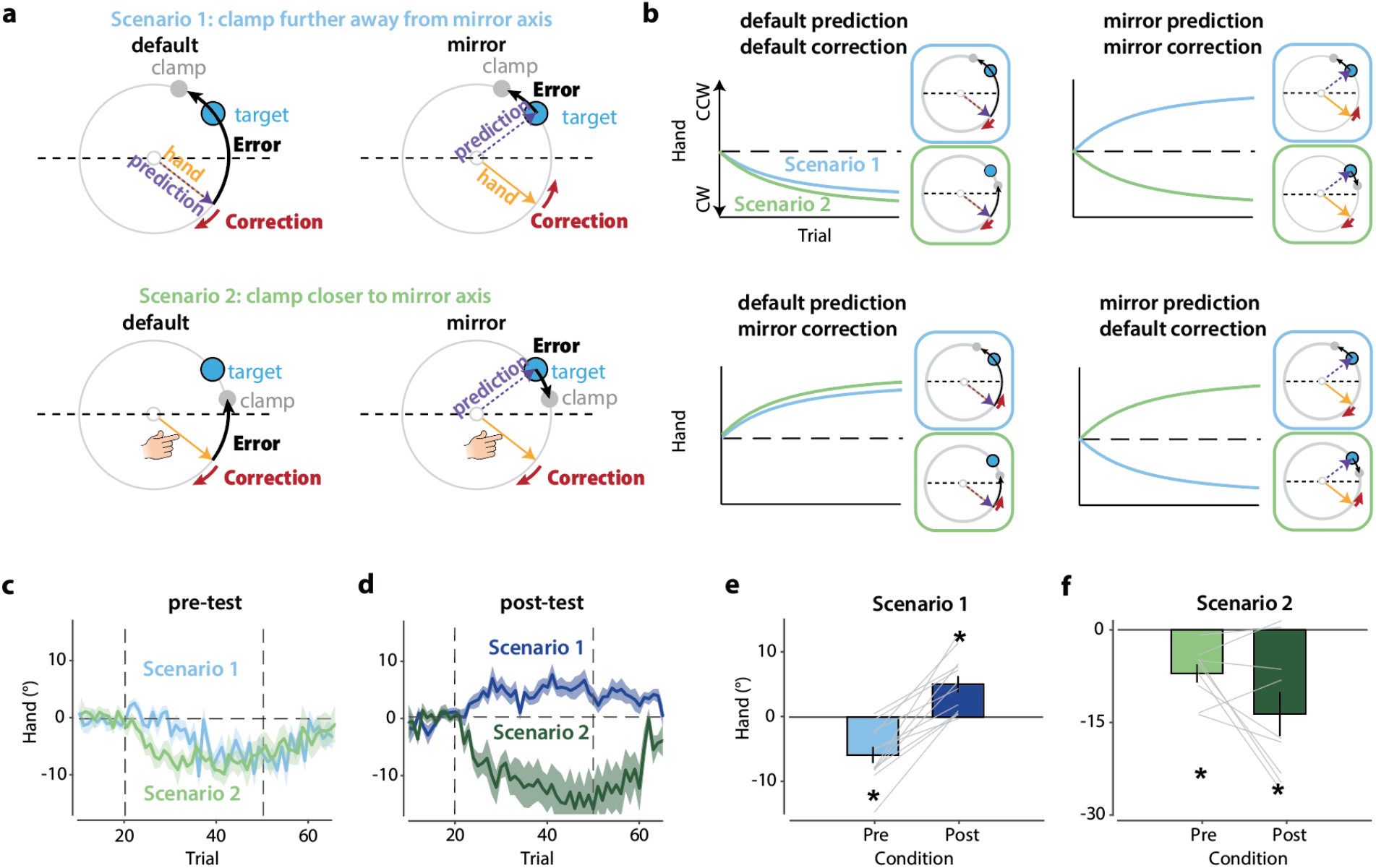
Implicit recalibration is properly reconfigured after prolonged mirror training. (a) Sensory prediction (purple dashed arrow) and error correction (red arrow) for implicit recalibration under the default and mirror mappings. A default-map operator predicts the cursor moves with the hand, whereas the mirror operator predicts a cursor trajectory reflected across the x-axis. The sensory prediction error (SPE) is the angular difference between the cursor feedback and this prediction (black arrow). The appropriate correction is opposite to the SPE under the default mapping but in the same direction under the mirror mapping. Two scenarios diagnose the operator after training. In scenario 1, the clamp is positioned farther from the mirror axis than the target, so the SPE points the same way under both mappings, but the correction diverges. In scenario 2, the clamp is closer to the axis than the target, so the SPE is opposite for the two mappings, but the correction is the same. (b) Predicted learning functions of implicit recalibration under four models that differ in whether the sensory prediction and motor correction are based on the default or mirror-reversed sensorimotor map. Blue and green traces show model predictions for scenarios 1 and 2, respectively. Panels to the right of the learning function duplicate the illustration in (a), but for each specific prediction-correction mapping combination. (c) Time courses of hand angle during the pre-test probe. The results are consistent with the model in which both the sensory prediction and error correction policy are computed within the default mapping. (d) Time courses of hand angle during the post-test probe after three weeks of training. The results are consistent with the model in which both the sensory prediction and error correction are computed within the mirror mapping (see panel b). (e–f) Bar graphs of hand angle averaged over the second half of the clamp phase in the pre-and post-test probes. The direction of the error correction reverses in scenario 1 (e) but not in scenario 2 (f), consistent with the model in which both the sensory prediction and error correction are computed within the mirror mapping. *, p < 0.005.

Crossing these two computations yields four possibilities following training. The sensory prediction could be based on either the default or mirror-reversed mapping, and the error correction policy could be based on either the default or mirror-reversed mapping. Importantly, these four possibilities yield unique behavioral predictions in two clamped feedback scenarios. In scenario 1, the clamp is positioned farther from the mirror axis (Fig 2a, upper), while in scenario 2, the clamp is positioned closer to the mirror axis (Fig 2a, lower). If prediction and correction are both based on the default mapping, the adaptive response is the same under both scenarios (Fig 2b, upper-left). However, if implicit recalibration is reconfigured to the mirror environment for both prediction and correction, the adaptive responses will be opposite in scenarios 1 and 2 (Fig 2b, upper-right). It is also possible that implicit recalibration is only reconfigured for either prediction or correction. Should this occur, the adaptive responses will be flipped relative to those predicted for full reconfiguration (Fig 2b, lower). Given the analytic value of the two scenarios, we examined the participants’ response to scenario 1 (n = 12) and a subset in scenario 2 (n = 9), prior to and after extended training in the mirror-reversed environment.

We observed clear reconfiguration of implicit recalibration after extended mirror-reversal training in Exp 1. At baseline, the participants’ response to clamped feedback under a mirror reversal matched the simulations in which the system operated on the default map (Fig 2c). However, after prolonged training, the behavior best matched the simulations in which both the predictions and error correction policy were based on a mirror-reversed map (Fig 2d). The consequence of this full reconfiguration is best highlighted in scenario 1, where the predicted direction of the correction is opposite for the default and mirror maps. Here we see a reversal from baseline to post-training in the shift in hand direction due to implicit recalibration, an effect evident in the behavior of each participant (Fig 2e, t(11) = 7.4, p < 0.001, ηp² = 0.83, BF₁₀ = 1630). Reconfiguration of implicit recalibration emerged slowly over the course of training. At the end of the first week, there was, on average, no change in heading in response to the clamp, and a clear reversal was observed after week 3 (Fig S1).

In contrast, under scenario 2, the predicted response direction remains the same if the maps for both prediction and correction have been reconfigured. This prediction matches the data (Fig 2f, t(8) = 1.9, p = 0.098, ηp² = 0.31, BF₁₀ = 1.1). It should be noted that the magnitude of adaptation in scenario 2 tended to increase after training, likely because the perceived error size changed due to the new predictions (Fig 2a, lower).

As an alternative probe of implicit recalibration, we applied hand-contingent perturbations that varied randomly in size and direction from trial to trial. Consistent with the results in response to clamped feedback, trial-by-trial corrections became properly scaled in response to the perturbation under the mirror map after training (Supplementary Results 1, Fig S2). In summary, using both non-contingent and contingent perturbations, Exp 1 provides compelling evidence that implicit recalibration can be reconfigured to operate under a mirror map after extended training. We next asked what specific aspects of training are necessary for this reconfiguration to occur.

### Training on center-out reaching is sufficient to reconfigure implicit recalibration

The results of Exp 1 demonstrate that, with sufficient training, implicit recalibration can be reconfigured to correct movement errors in an appropriate manner for a given sensorimotor map. Given that this flexibility is at odds with the results from previous mirror-reversal studies^6–8,11,16,17^, Exp 2 was designed to assess which feature or features of our training procedure account for the difference. One possible reason is that our training protocol included tasks that engage online feedback control, a feature largely absent in prior studies in which training was restricted to center-out reaching. It is possible that feedback-based online corrections may help teach the feedforward system to learn the appropriate error correction policy^20,26,27^. A secondary analysis of the data from Exp 1 revealed that feedback control improved following our extended training. As part of the post-test battery, we included a target-jump task and found that, when the target was displaced mid-reach, participants became more efficient at making rapid, online corrections under the mirror reversal (Supplementary Results 2, Fig S3).

To examine the role of feedback-based training, we trained two new groups of participants with a mirror-reversed mapping using different task sets. For one group, the training was limited to center-out reaching (i.e., feedforward-biased training); for the other group, training was limited to continuous tracing (i.e., feedback-biased training). Participants improved on their respective task across the 7-day training period (Fig 3a). As in Exp 1, we focused on how the participants responded to clamped feedback under a mirror mapping using the stimulus configurations of scenario 1 since these provide the most discriminative test.

**Figure 3.**
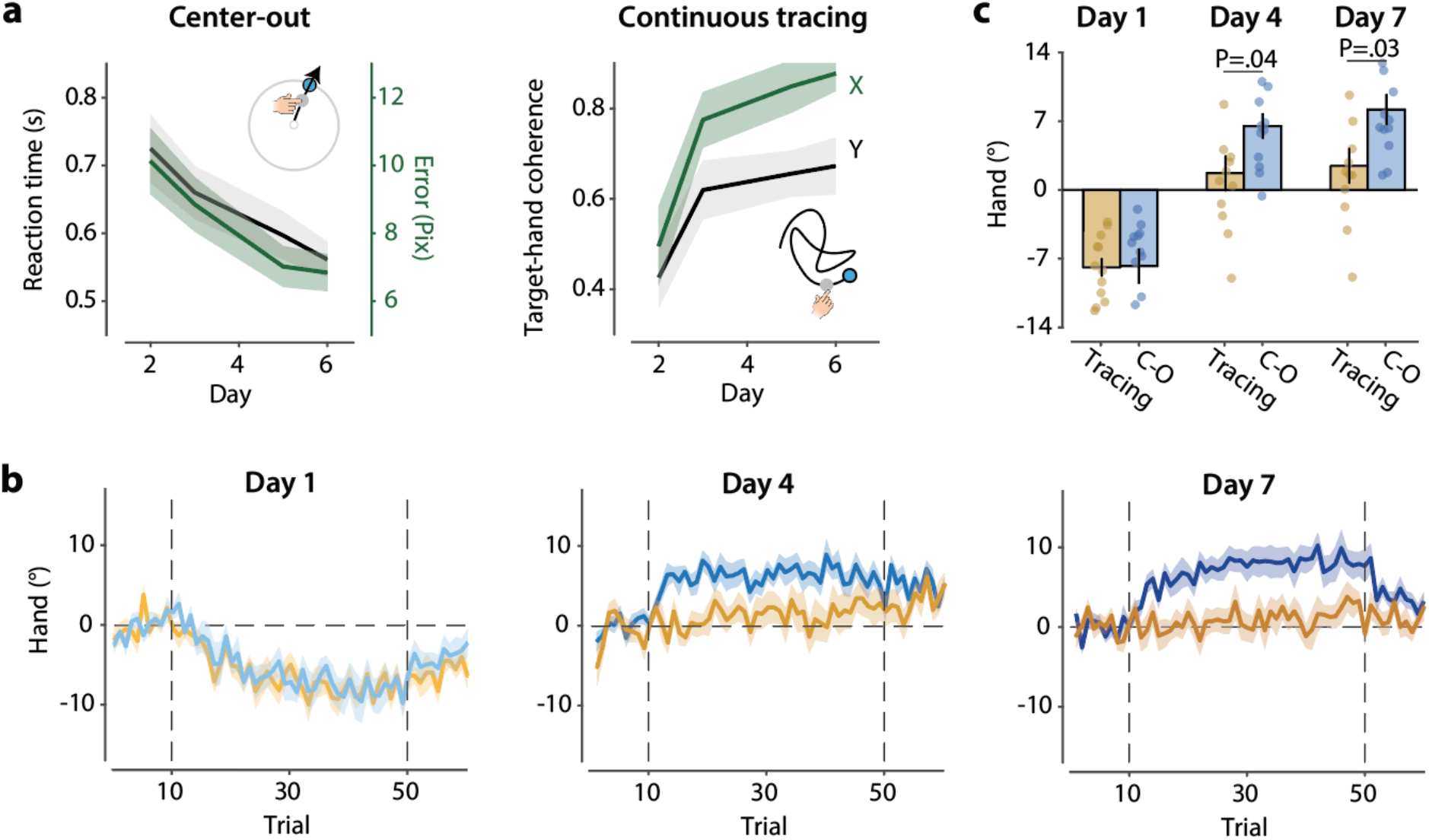
Center-out reaching task is more effective than continuous tracing in reconfiguring implicit recalibration. (a) Learning curves over one week of training for two groups, trained exclusively on either the center-out reaching task (top) or the continuous tracing task (bottom). Both groups improved on their respective task across training days. Linear mixed-effects models revealed significant improvements in reaction time and reach error for the center-out group, and in target–hand coherence along both the non-perturbed (X) and mirror-reversed (Y) axes for the tracing group (all |z| > 4, p < 0.001). (b) Time course of implicit recalibration measured with clamped feedback in the pre-test (day 1), mid-test (day 4), and post-test (day 7). We only tested the condition in which the clamp is displaced away from the mirror axis since the corrective response will reverse direction to this perturbation if implicit recalibration is based on a mirror-reversed map (see Fig 2a–b, scenario 1). In the pre-test, both groups showed adaptation aligned with the default map. Only the group trained on the center-out task showed a reversal following training. (c) Mean hand angle during the second half of the clamp block for each probe day.

While both groups showed that recalibration was based on the default mapping in the pre-test (center-out t(11) = −4.3, p = 0.001, ηp² = 0.63, BF₁₀ = 35; tracing t(11) = −8.4, p < 0.001, ηp² = 0.86, BF₁₀ = 4.5×10³), they sharply diverged over the course of training (Fig 3b–c). Surprisingly, we observed a clear reconfiguration of implicit recalibration in the center-out group after just 4 days of training (t(11) = 5.0, p < 0.001, ηp² = 0.70, BF₁₀ = 89). This reconfiguration persisted through the end of training (day 7) and reached levels comparable to those observed after three weeks of training from Exp 1 (t(11) = 5.1, p < 0.001, ηp² = 0.71, BF₁₀ = 105). In contrast, the response to the perturbation for the continuous tracing group had flattened out by day 4 (t(11) = 0.94, p = 0.37, ηp² = 0.074, BF₁₀ = 0.42) and remained so on day 7 (t(11) = 1.3, p = 0.21, ηp² = 0.14, BF₁₀ = 0.59), providing no evidence of appropriate responses for the mirror map. While the participants’ behavior did change compared to the pre-test on day 1, it is unclear if the flattened functions reflect a slow transition to the mirror map, a mixture of mappings, or suppression of the adaptive response.

Additional analyses further confirm that the difference in implicit recalibration between training groups is not due to differences in online-control ability (Supplementary Results 3, Fig S4). As such, online control is not required for the emergence of implicit recalibration based on a novel sensorimotor map. Rather, training based on feedforward planning alone appears to be sufficient. It remains puzzling why the degree of reconfiguration of implicit recalibration reported in prior mirror-reversal studies was so much less than what we have observed here as it cannot be attributed to training duration or differences in task set. We return to this issue in the Discussion, pointing to methodological differences that may account for limitations in previous studies.

### Flexibly switching between the default and mirror maps

Having shown that implicit recalibration can be reconfigured to operate under a novel sensorimotor map, we next asked how this new state would impact performance when the mirror reversal was removed. If implicit recalibration is tied to a single mapping, then we should observe interference: corrections would initially be maladaptive and require some degree of relearning. Alternatively, it may be that, once learned, implicit recalibration can be flexibly reconfigured to be appropriate for a given mapping, a form of cue-dependent learning.

To evaluate these hypotheses, we conducted two tests on the trained participants (n = 12) from Exp 1. After the post-test, participants were tested on a block of trials with the reversal removed (i.e., feedback based on the default map; Fig 1h). Before this test, participants were explicitly told that the mirror manipulation had been turned off and that they should reach directly to the target. Interestingly, they showed typical implicit adaptation, indicating that implicit recalibration was operating on the default map (Fig 4a; t(11) = −6.0, p < 0.001, ηp² = 0.77, BF₁₀ = 317). Thus, extended exposure to a novel sensorimotor map does not lead to an overwrite of a system tuned to the default map but instead builds a new implicit operator, tuned to the mirror-reversed context.

**Figure 4.**
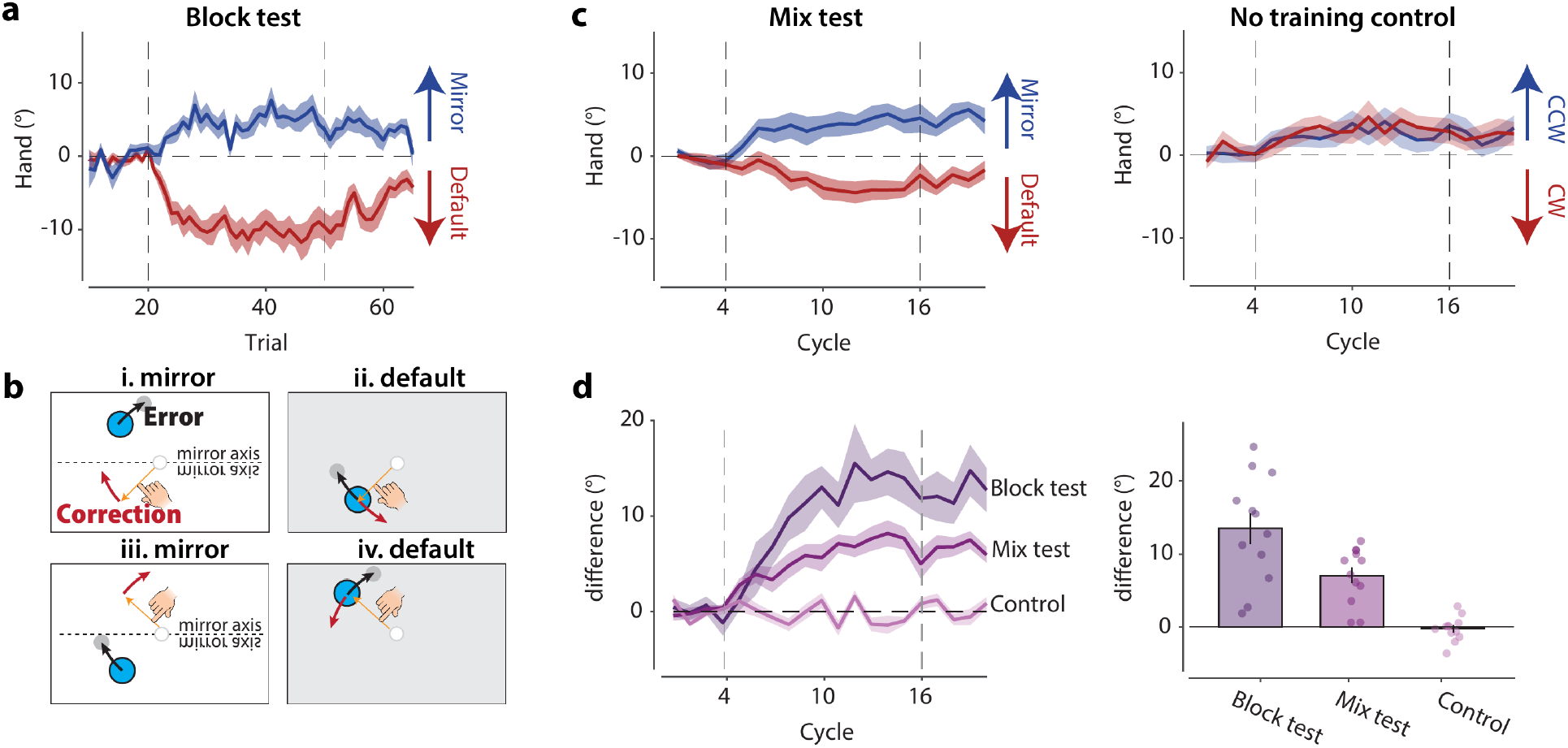
Low interference between mirror and default mappings. (a) Learning curves for the two mappings measured in independent post-test sessions (block test). (b) Illustration of the mixed test assessing trial-by-trial flexibility of implicit recalibration. Four conditions, created by the combination of 2 mappings and 2 targets, were randomly intermixed. The mapping for a given trial was indicated by the background color. The perturbation was selected such that movements with matched motor commands (i & ii; iii & iv) would produce changes in hand angle in opposite directions if implicit recalibration operated on the appropriate map. (c) Time course of reaching direction in the mixed test (left) compared with a naïve control group (right), in which participants adapted to clamps in opposite directions (CCW/CW) cued by background color under the default mapping. Participants in the mixed test simultaneously adapted in two directions across the two mappings, whereas control participants could not separate the opposite clamps on the basis of background color. (d) Time course of the difference in reaching direction between the two mappings (left) and the bar graph of this difference averaged over the second half of the clamp phase (right), for the block test, mixed test, and control group.

The coexistence of two implicit operators raises a further question: can the implicit recalibration system switch between two mappings on a trial-by-trial basis? Prior work has shown that implicit memories can be separated when opposing perturbations are linked to distinct contextual cues, such as visual motion or follow-through movements^28,29^. However, when learning is constrained to the implicit system, these effects are typically very small (< 1°)^30,31^. We therefore conducted a mixed test (a dual-adaptation design), asking whether the sensorimotor map can serve as a contextual cue that separates the operation of implicit recalibration. The same participants from Exp 1 were given an additional week of training in which the default and mirror mappings alternated on a trial-by-trial basis, cued by the background color before each trial. To ensure that movement and aiming directions overlapped across the two mappings, we used target pairs positioned symmetrically about the mirror axis (Fig S5). Critically, we selected clamps that would be expected to drive adaptation in opposite directions under the default and mirror maps (see Fig 4b).

Participants showed simultaneous adaptation in opposite directions depending on the cue, with a clear separation of the two learning curves (Fig 4c; t(11) = 6.4, p < 0.001, ηp² = 0.79, BF₁₀ = 502). Thus, the adaptation system was sufficiently flexible to switch on a trial-by-trial basis with minimal interference. To confirm that this separation came from the learned mappings rather than the visual cues alone, a naïve group with no mirror training was exposed to the same visual cues, one color associated with a CW rotation and the other color associated with a CCW rotation. These participants showed no separation between the responses to the two cues (Fig 4d; t(11) = −0.54, p = 0.60, ηp² = 0.026, BF₁₀ = 0.33) and the comparison between the trained and naïve groups was highly significant (Fig 4d; Welch’s t(16.2) = 8.0, p < 0.001, ηp² = 0.80, BF₁₀ = 1.3×10⁵). As such, the results show that once a new mapping is learned, participants can effectively maintain and switch between two operators that require opposing error correction policies. Moreover, this degree of context-dependent flexibility is much greater in magnitude than that observed with other context-dependent implicit measures^30,31^.

### Implicit recalibration can also be reconfigured for a rotational map

The preceding results led us to revisit a well-studied mapping, visuomotor rotation, asking whether a new operator would be learned in this case. Unlike mirror reversals, where the default map leads to a correction in the wrong direction, the change in hand angle to a rotational error is in the appropriate direction. However, various lines of evidence suggest that implicit recalibration saturates in terms of how it can be used to modify the default map, with estimates of the upper bound falling around 20–30°^22,32,33^.

Given this limit, learning to compensate for a larger rotation has been hypothesized to require an additional explicit re-aiming component. For example, compensating for a 90° rotation might entail a combination of 20° of implicit recalibration plus an explicit change in aiming of 70°.

While this dual-process model is sufficient to compensate for the rotation, relying on aiming strategies can be costly, requiring substantial cognitive effort^34–36^. Our mirror-reversal results suggest a solution: prolonged practice with a large rotation could enable the system to construct a new operator that acts directly on the rotated mapping, with implicit recalibration reconfigured to this new map.

To examine this idea, participants in Exp 3 were trained on a 90° visuomotor rotation for two weeks using a center-out reaching task (Fig 5b). Performance, measured in terms of hand angle error or reaction time, improved across the training period (Fig 5c). To probe whether implicit recalibration was reconfigured over the course of training, we applied a clamped perturbation on top of the rotation during pre-and post-test assessments. The predicted direction of implicit recalibration differs depending on whether implicit recalibration is operating on the default map or has been reconfigured to operate on the rotated map (Fig 5a). Critically, the policy for the motor correction to a given error is the same for both mappings—correct the movement in the opposite direction of the error. As such, a reversal in the direction of the correction would mean that there is a change in the sensory prediction.

**Figure 5.**
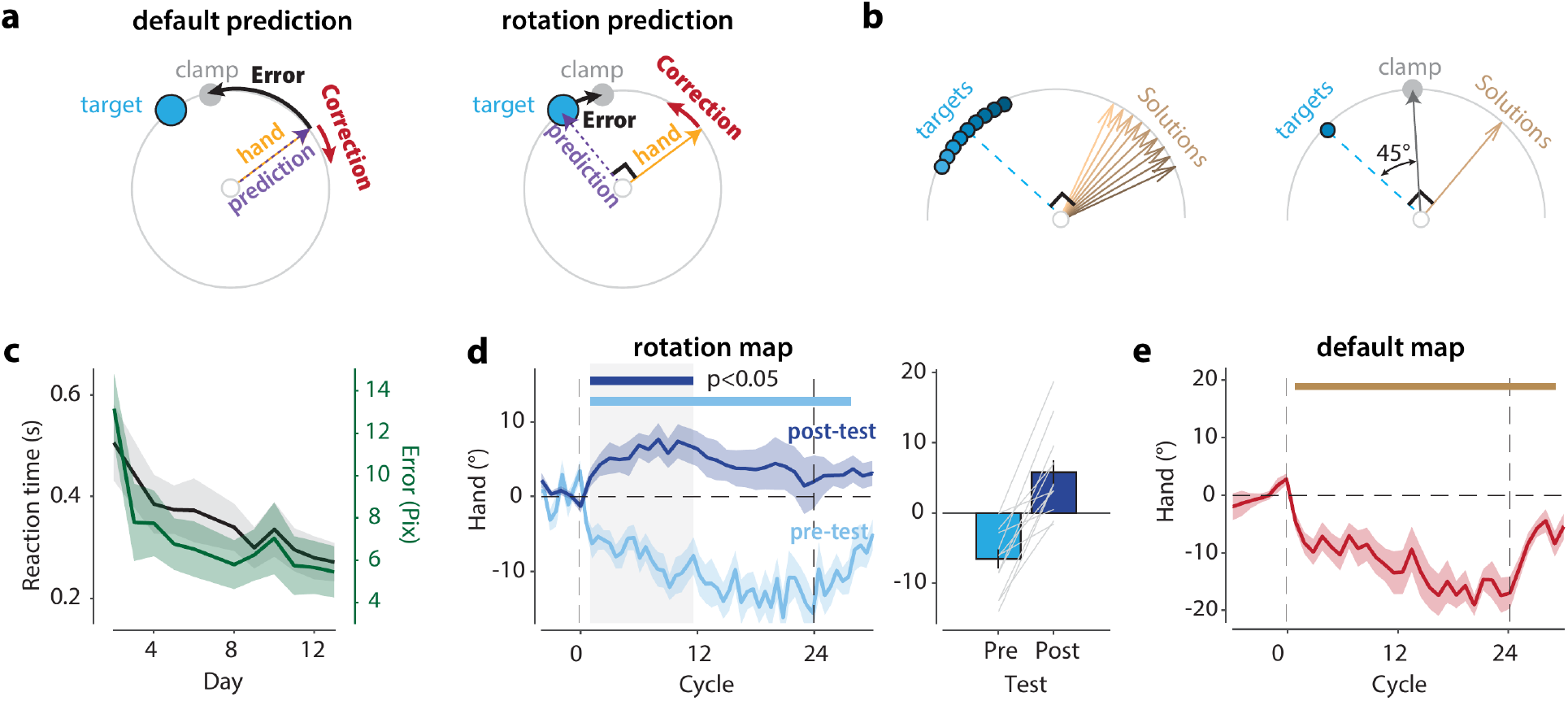
Implicit recalibration is reconfigured during prolonged training with a visuomotor rotation. (a) Predicted direction of correction to a clamp perturbation imposed on a 90° counterclockwise rotation. Left: Schematic showing that the response of untrained participants is in the same direction as the clamped perturbation, in accord with the hypothesis that the error is defined as the difference between the predicted feedback (expected hand position) and the cursor (default mapping)^15,37^. Right: If, with training, the mapping is reconfigured to be based on the rotation, the prediction would now be at the target and the change in hand angle would be in the opposite direction of the clamped perturbation. (b) During training, participants practiced a rotation using targets spanning 115–155°, with the corresponding solutions spanning 25–65°. As shown for one target location, the clamp is positioned 45° CW from the target. All nine trained targets were tested in the post-test, though only the center target is shown here for illustration. (c) Performance improved on measures of reaction time and accuracy over the two-week training period. Linear mixed model: both zs < −8.5, ps < 0.001. (d) Left: Time course of hand angle over trials when clamped feedback was imposed on top of the 90° rotation. At pre-test, implicit recalibration operated on the default map; at post-test, the system operated on the rotational map. Target configurations were slightly different between pre-and post-test (see Methods). The thick bars indicate epochs during which the hand angle differed significantly from zero. Right: Mean hand angle during the overlapping significant period of the pre-and post-test time courses (gray area). (e) Time course of hand angle in response to clamped feedback when the 90° rotation was eliminated and participants were instructed to reach directly to the target. There was no indication of interference in using the default map after the rotational training.

For the pre-training probe, participants were explicitly informed of the rotated mapping and given the appropriate aiming strategy. This allowed them to fully compensate for the rotation within a brief familiarization block (5 trials per target). When the clamped perturbation was superimposed on the 90° rotation, implicit recalibration aligned with the default map (Fig 5d; pre-test vs. 0: t(11) = −4.75, p < 0.001, ηp² = 0.67, BF₁₀ = 62), producing corrections that were inappropriate given the rotation (i.e., changes that would increase movement error). However, after two weeks of training, the direction of the responses to the clamped perturbation was reversed, now producing changes in hand angle that were appropriate for a rotated map (Fig 5d; pre-test vs. post-test: t(11) = 5.0, p < 0.001, ηp² = 0.69, BF₁₀ = 85; post-test vs. 0: t(11) = 2.3, p = 0.041, ηp² = 0.33, BF₁₀ = 1.9). Thus, a new operator had been created, one based on a sensorimotor map that incorporated the rotation, leading implicit recalibration to produce responses appropriate for the error.

Given that the error correction policy is the same at pre-and post-training, we assessed if learning the rotated mapping interfered with implicit recalibration under the default mapping in a subset of participants (n = 10). Similar to Exp 1, we removed the 90° rotation and instructed participants to reach directly to the target. When the clamp perturbation was introduced, participants showed a learning curve in the direction appropriate for the default mapping; thus, we found no evidence of interference (Fig 5e; default-map adaptation vs. 0: t(9) = 5.9, p < 0.001, ηp² = 0.79, BF₁₀ = 140). The similarity of the results for training with either mirror-reversed or large rotations suggests a general principle: implicit recalibration can be reconfigured to operate under novel sensorimotor maps in a flexible manner with minimal interference between configurations.

## Discussion

A hallmark of skilled movement is the ability to adjust behavior fluidly in response to unexpected events or changing conditions^1,2^. This ability is thought to be supported by implicit sensorimotor recalibration, an automatic process that compensates for errors and keeps the sensorimotor map accurate^6,7,14,15^. Implicit recalibration is well characterized in simple reaching tasks, but it remains uncertain whether it also supports everyday skills in which the mapping between motor commands and their sensory consequences is not always intuitive, such as when backing up a car, trimming a sail, or cutting one’s own hair in the mirror. Whereas our everyday experience suggests that we routinely and effortlessly recalibrate skilled movements, laboratory studies of implicit recalibration have highlighted how this system appears to be remarkably rigid, operating within a fixed, intuitive reference frame that cannot be reconfigured when the mapping changes^6,22,37–39^. Here, we reconcile this apparent paradox. With prolonged practice on a new mapping, implicit recalibration reversed direction to become appropriate for that mapping, generating context-specific corrections and revealing a flexibility previously thought impossible for this system.

The prevailing characterization of implicit recalibration, built largely on center-out reaching, holds that the system is encapsulated and insensitive to the learner’s explicit knowledge of the task: it corrects errors according to a default operator even when doing so is maladaptive^6,37,38^. Consistent with this view, previous studies have repeatedly found that implicit recalibration under a mirror reversal remains locked to the default mapping and thus generates ‘corrective’ responses that actually increase error, a pattern that persists even after a week of training^6–8,11,16,17^. This raised the question of whether implicit recalibration can ever be reconfigured for a novel mapping, or whether error correction for these mappings must be supported by some system other than sensorimotor recalibration.

Our results mandate a fresh perspective on these questions. We observed a lack of flexibility for implicit recalibration early in training. However, with extended and appropriate practice, implicit recalibration comes to compensate properly for errors under a mirror reversal. We identified the two core computations required to produce mapping-specific responses and found that both were reconfigured. First, the sensory prediction came to anticipate the mirrored consequence of the movement. Second, the error-correction policy came to drive the hand in a direction appropriate for the mirror-reversed environment.

Our results also revise a common assumption about the role of online feedback control in skill learning: namely, that online error correction is a key ingredient for the implicit system to learn a new error correction policy. Indeed, when embarking on this project, we thought the absence of training with online feedback might explain why earlier studies failed to find successful reconfiguration of implicit recalibration under a mirror reversal^16,18–21^. But this proved not to be the case. Training on a mirror reversal with center-out reaching alone was sufficient to reverse implicit recalibration. In contrast, training for a similar amount of time with continuous tracing alone resulted in suppression of the maladaptive response but no reversal. We do note that online control became more efficient with practice in both groups (Supplementary Results 3). Thus, the capacity for online control is not the driver of the reconfiguration of implicit recalibration, and instead appears to reflect the development of a separate skill.

Why, then, did earlier studies not observe reconfiguration of implicit recalibration^6,7,9,40^? Our results argue against the most obvious explanations. It is not simply a matter of training duration: a clear reversal emerged after only four days of center-out practice in Exp 2, well within the range of previous multi-day studies^6,7,40^. Nor is it a matter of training variety, as the single, typical center-out task was sufficient.

We propose several possible explanations for why the past studies failed to observe the reconfiguration. First, there are substantial methodological differences in how the stimuli were presented and the way the responses were made between the current and previous work. We used a laptop-based system in which a mouse was moved in the horizontal plane and the visual display was vertical. As such, movement space and cursor space were physically dissociated. Under this condition, the mirror map inverts a mapping that is non-veridical yet highly familiar, the one with which we operate a standard mouse. In previous lab-based studies^6,7,9,40^, the cursor was projected on a horizontal screen and arm movements were made on a table/tablet that was physically aligned with the cursor under the default map. This spatial alignment may promote stronger integration of visual and proprioceptive feedback that is harder to override^41,42^, such that reconfiguring implicit recalibration may require much longer training.

Second, the difference may lie in how implicit recalibration was probed. We assayed implicit recalibration in the current study by imposing an additional perturbation on top of the mirror map. In previous studies, the implicit component was measured from baseline reaches to the target under the mirror reversal^6,7,40^. We suspect that this procedure measures how participants adjust their control policy to the mirror environment rather than the operation of implicit recalibration. By applying an additional perturbation in the current experiments, we directly probed implicit recalibration. Third, in some of the previous studies^6,7^, implicit recalibration was inferred by having participants report their aiming direction on each trial. It is possible that emphasizing an explicit aiming strategy may lead the participants to attempt to solve the task through explicit cognitive substitution^43^, and that this may interfere with the ability of the implicit system to build a new operator.

Notably, the reconfiguration of implicit recalibration was not restricted to mirror reversal but was also observed when the perturbation required responding to a large visuomotor rotation, one that well exceeds the capacity for implicit recalibration^11,22^. The reconfiguration to a new mapping associated with the large rotation speaks to another puzzling phenomenon in the adaptation literature. Even when participants are given an explicit aiming strategy to nullify a large rotation, they continue to adapt implicitly during training, with the hand angle gradually drifting away from the target. In essence, performance deteriorates with practice^15,37^. This maladaptive behavior arises because implicit recalibration remains anchored to the default map even when that mapping is no longer appropriate.

Although we observed exactly this maladaptation in the pre-test, two weeks of training with a large rotation resulted in implicit recalibration becoming reconfigured to the rotated map, correcting errors in an appropriate manner. In this sense, this phenomenon does not reflect a fixed property of the implicit system but a situation in which the explicit and implicit systems do not coordinate in the short term^9,11^. With extended practice, the system constructed a new operator that operates directly on the rotated mapping. As with mirror training, the new operator eliminated the need for the aiming strategy; participants came to aim at the target itself rather than at a rotated surrogate and implicit recalibration generated an appropriate response.

A striking feature of our results is the minimal interference between the mirror and default maps. A color cue was sufficient to allow participants to switch between two operators, revealing that the implicit system can maintain and select amongst parallel, context-specific operators. Similar effects have been observed in many studies in which the sensorimotor map is the same but the direction of the perturbation is reversed for the two cues^28,44,45^. The concept of a neural manifold, the low-dimensional space to describe the activity of neural populations^46,47^, offers one way to conceptualize this flexibility. Implicit recalibration under a default mapping can be viewed as corrections within an existing manifold^48^. These could be readily achieved without requiring any change in the control policy. In contrast, acquiring a new skill might require the establishment of a new manifold, a process that requires extended training to build the new operator^49,50^. Once a new manifold is established, the system can again exploit it for fast adaptation, now appropriate to the new mapping.

How does the cerebellum, the structure most closely associated with implicit recalibration, fit into this picture? Classic models propose that the cerebellum learns multiple paired forward (predictor) and inverse (operator) models and selects the pair appropriate to the current context through a responsibility signal^24,51^. On this account, mirror training builds a new forward–inverse pair dedicated to the mirror context while leaving the default pair intact. However, recent work suggests that the cerebellum contributes to the control policy itself, acting as an operator rather than a pure forward-model predictor^8,52,53^. On this view, the cerebellum may continue to generate corrections in default coordinates, while the cortex learns the new sensory prediction and re-maps the cerebellar output so that it acts in the correct direction under the new mapping. To distinguish the relative contributions of the cerebellum and cortex, future behavioral or physiological work should examine whether the predictive and corrective components are jointly reconfigured, or whether this process occurs with some degree of independence. Answering these questions will shed light on how the reconfigured map is represented within the sensorimotor hierarchy.

For over a century, anecdotal reports have described how the ability to move in complex environments is possible after extended experience with inverted or mirror-reversed vision^12^. These reports have sat uneasily with laboratory evidence indicating that implicit recalibration is rigidly bound to a default sensorimotor map^6,7,14,15^. Here we resolve this puzzle with clear behavioral evidence that, with sufficient training, implicit recalibration can be rebuilt to serve a new mapping, whether a mirror reversal or a large rotation, and the resulting operators coexist and can be selected by context. The implicit system, long viewed as the inflexible component of motor learning, is thus reshaped by experience, providing a foundation for skills that depart from the intuitive geometry of the body.

## Methods

### Participants

A total of 77 participants (47 women, age: 23.4 ± 4.2 yr) were recruited. All participants were right-handed and possessed normal or corrected-to-normal visual acuity. The participants were financially compensated for their time. The protocol was approved by the Institutional Review Board at the University of California, Berkeley (approval number: 2016-02-8439). Informed consent was obtained from all participants.

### Apparatus

The tasks were administered on participants’ personal computers. Experiments were implemented in JavaScript and run in the Google Chrome browser, with data stored in Google Firebase^54^. Participants were instructed to sit still. The laptop was placed ∼50 cm from the body midline, and a mouse pad was placed to the right of the laptop. For most participants, movements were made using the right hand to move a vertical mouse on the mouse pad; the exception was three participants in Exp 1, who used a digitizing pen on an external tablet. Participants were instructed to replicate this setup at home, which we confirmed over Zoom. A white circular cursor (radius: 0.85% of screen height) provided real-time feedback of hand position. The system enforced a fixed gain, such that cursor displacement was a constant function of hand displacement, independent of movement speed.

### Behavioral tasks

Participants were trained to move under two types of novel mappings. In Exps 1 and 2, we imposed a mirror-reversal mapping in which the vertical position of the feedback cursor was inverted across the horizontal axis. Thus, an upward hand movement produced a downward cursor movement. The horizontal position of the feedback cursor was unchanged. In Exp 3, we imposed a visuomotor rotation mapping in which the position of the feedback cursor was rotated 90° counterclockwise about the start position relative to the actual hand position (contingent feedback).

To increase training variability, we used three tasks during the training phase: continuous tracing, point-to-point reaching, and center-out reaching. These were selected to provide training with both online control and feedforward planning. We probed implicit recalibration with two approaches: non-contingent, task-irrelevant error clamps and random, trial-by-trial perturbations of contingent feedback. To assess the impact of training on online control, we used a target-jump task. Each task is detailed below. In describing the target position and perturbations in angular coordinates, 0° is defined as pointing east. Positive angles indicate a counterclockwise rotation; negative angles indicate a clockwise rotation.

### Center-out reaching

Center-out reaching emphasizes feedforward planning. Each trial began with the participant positioning the cursor within the starting circle at the center of the screen and maintaining this position for 500 ms. A blue circle, the target (radius: 1.8% of screen height), appeared at a fixed radial distance of 30% of screen height, with its angular position randomly selected from a uniform distribution of 60 possible locations (0–354° in 6° increments). Each target location was repeated twice in a row to encourage error correction, resulting in 120 trials per day. Participants were instructed to reach to the target as quickly and accurately as possible, slicing through the target before returning to the start location. If movement time (defined as the interval from movement onset to the time when movement amplitude reached the target distance) exceeded 500 ms, the message “Too Slow” was displayed for 500 ms. The perturbed cursor (mirror-reversed or rotated) was presented only during the outward reach. Once the movement amplitude reached the target distance, the target and cursor disappeared and the starting circle reappeared. To aid the participant in finding the center position, the cursor, veridically aligned with hand position, reappeared when the hand was within 5% of screen height from the start position.

### Continuous tracing

Continuous tracing emphasizes online control, and we adopted the exact design from previous work^11^. At the start of each trial, participants moved the cursor to the starting circle and maintained this position for 500 ms. The target then moved away from the center along a pseudo-random trajectory and participants needed to trace it as precisely as they could. The target followed a two-dimensional path defined as a sum of sinusoids along a given axis at time t: r(t) = Σᵢ [aᵢ · sin(2π·ωᵢ·t + ϕᵢ)]. For the x-axis, amplitude (a/pix), frequency (ω/min^-1^), and phase (ϕ) were [100, 100, 100, 50, 50, 30, 30], [7, 16, 34, 49, 67, 91, 121], and [0, π/4, π/2, 3π/4, π, 5π/4, 3π/2], respectively. For the y-axis, these parameters were [100, 100, 50, 50, 50, 30, 30], [7, 19, 40, 58, 85, 109, 127], and [0, π/3, 2π/3, π, 4π/3, 5π/3, 2π]. Parameters were chosen so that larger amplitudes and velocities occurred primarily at lower frequencies (0.1–0.65 Hz), with minimal spectral overlap between axes. Participants were instructed to track the moving target as accurately as possible using the cursor. The target disappeared after 1 minute of tracing, at which point the starting circle reappeared with the participant instructed to return to the starting location to initiate the next trial. Participants completed 5 one-min trials in each training session in Exp 1 and 12 trials in Exp 2.

### Point-to-point reaching

This task was selected as a hybrid of continuous tracing and center-out reaching, requiring both feedforward planning and feedback control. Each trial began with the participant moving the cursor to the start location and maintaining this position for 500 ms. A target (radius: 1% of screen height) then appeared 150 pixels from the start position with its angular position selected at random. The participant was instructed to move the cursor to the target as quickly and accurately as possible, with the requirement that the cursor land on the target. Thus, corrective movements were required if the cursor missed the target. Once the target was intersected, a new target immediately appeared 150 pixels from the previous target with the angular position selected at random. Trials were grouped into cycles of six sequential targets. Once the final target was reached, the start circle reappeared, signaling the start of the next cycle. Each training session included 240 trials (40 cycles).

### Clamp adaptation task

For probes with clamped feedback, we used a variant of center-out reaching^5^. For this perturbation, the radial position of the cursor tracked the hand, but the angular position followed a fixed angular trajectory relative to the target. Thus, the angular position was independent of the participant’s actual movement direction (non-contingent feedback). Participants were informed about the nature of the feedback and instructed to ignore the cursor and reach directly to the target. As shown in many studies with a default sensorimotor map, despite these instructions, participants gradually shift their reach direction in the opposite direction of the clamped feedback. This response is automatic, reflecting the operation of implicit recalibration attempting to nullify the visual error.

For each clamp session under mirror reversal, three targets were used. For one set of targets, the locations were 45°, 135°, and 180°; for the other set, the locations were 315°, 225°, and 0°. To limit generalization from exposure to the clamps, the two target sets were employed in separate sessions, and participants performed both in pre-and post-test assessments. By varying the direction of the clamped feedback, we created two scenarios (see Fig 2a). Scenario 1 used clamps that would result in responses that would differ in direction if implicit recalibration operated on the default or mirror map. For this scenario, the 45° and 225° targets were paired with a 30° clamp, and the 315° and 135° targets with a −30° clamp. Scenario 2 used clamps that would result in responses in the same direction, regardless of whether implicit recalibration operated on the default or mirror map. Here the 45° and 225° targets were paired with a −30° clamp, and the 315° and 135° targets with a 30° clamp. We also included two targets on the mirror axis (180° and 0°) and always associated them with a 30° clamp. However, we do not report the results for these targets as they do not require learning a new map and it is unclear how to define the proper error correction policy. Each clamp session consisted of 5 cycles with contingent feedback (3 targets per cycle), followed by 15 baseline cycles without feedback, 30 cycles with clamped feedback, and 20 cycles without feedback. The first set of no-feedback cycles was included to establish baseline performance, and the last set of no-feedback cycles provided a probe of washout from the clamped feedback.

As a pre-training baseline, we measured implicit recalibration under the mirror reversal before extended training. Before this pre-test assessment, we provided explicit instructions to describe the mirror-reversed mapping and provided a verbal aiming strategy (e.g., aim at −45° to reach a target presented at +45°). Participants then completed a brief training block (5 trials per target) to confirm that they understood the perturbation and could successfully implement an appropriate aiming strategy. We then described the clamped feedback and instructed participants to ignore the cursor and simply aim their reaches according to the mirror rule. This same procedure for the clamped feedback probe was administered after each week of training.

To examine whether implicit recalibration can flexibly switch across mappings, targets were arranged in pairs positioned symmetrically relative to the mirror axis (Fig S5). By using symmetric targets, the same motor command for a given target was shared across the two mappings. For these participants, eight targets were used, forming four mirror-symmetric pairs, all positioned off the mirror axis (112°/248°, 116°/244°, 124°/236°, and 128°/232°); no target lay on the axis. The relevant mapping on each trial was signified by the background color; a black background indicated the mirror mapping and a gray background the default mapping. Each target appeared in both backgrounds, and the sign of the clamp offset reversed between them. Under the mirror mapping, the clamp was +30° for targets above the mirror axis (112°, 116°, 124°, 128°) and −15° for targets below it (232°, 236°, 244°, 248°). Under the default mapping, every sign was reversed (Fig 4b). As such, the adapted response would be in opposite directions for the mirror and default maps. Each session began with a block of no-feedback baseline trials (80 trials) followed by a block of clamped-feedback trials (480 trials), and ended with another no-feedback washout (80 trials). Within each block, trial order was randomized in successive cycles of 16 trials (8 targets × 2 mappings).

### Random perturbation

We also used a second probe of implicit recalibration, one that involved contingent cursor feedback. For this probe, the cursor was rotated on each trial by a random angle relative to the participant’s movement direction^33^. The direction (CW or CCW) and magnitude of the rotation (±5°, ±10°, or ±15°) varied randomly across trials. Participants were told that the feedback would change unpredictably and instructed to reach directly to the target, ignoring the feedback (similar to the instructions for the clamped feedback probe). Trials were organized into sets of seven that shared a common target, with a new target used for each set. The first six trials of a set presented each of the six perturbation conditions (±5°, ±10°, and ±15°) once, in random order; the seventh trial was unperturbed (0°) and provided veridical endpoint feedback. Each participant completed 90 sets (630 trials), with target directions randomly sampled from 0° to 357° in 3° steps.

### Target-jump task

We used a target-jump task to examine how training impacted online control^16^. The basic task was similar to the center-out task, with the participant instructed to reach quickly and accurately to a target. Approximately 100 ms after movement onset, the target was shifted to a new location on two-thirds of the trials, displaced by either +30° or −30°. On the other third of the trials, the target position was unchanged; thus, both the presence and direction of the displacement were unpredictable. The trial ended on no-jump trials when the cursor exceeded the target distance. For jump trials, the trial concluded once the cursor entered a circular zone (radius: 2% of screen height) centered on the new target location. The target appeared at one of six possible locations (0°, ±60°, ±120°, and 180°). The experiment began with a 36-trial warm-up with contingent online feedback and no target jumps, followed by 270 intermixed trials in which each target was presented 45 times (15 trials each for the +30° jump, −30° jump, and no-jump conditions), with their order randomized in cycles of 18 trials (6 targets × 3 jump types).

### Training procedure

#### Experiment 1: Long-term mirror-reversal training

To examine whether implicit recalibration can be reconfigured to a novel sensorimotor map, participants (n = 12) were trained on a mirror-reversal task for approximately one month. We describe the general experimental design here (Fig 1h), although individual participants followed slightly different training schedules (see Supplementary Methods).

Prior to training, we obtained baseline measures on the training tasks under a default visuomotor mapping (veridical feedback), as well as assessed pre-training behavior in response to clamped feedback under a mirror-reversed mapping. The latter consisted of two blocks, one with the scenario 1 clamps and one with the scenario 2 clamps.

For mirror training, each daily session included 5 continuous tracing trials, 240 point-to-point reaching trials, and 120 center-out reaching trials. These three tasks took approximately 20–30 minutes to complete. Participants trained for five consecutive days each week. On the sixth day, implicit recalibration was assessed using the clamp adaptation task (scenario 1 only). No testing was performed on the seventh day.

During the first three weeks, participants trained exclusively under the mirror-reversal mapping. A clamp test (scenario 1 only) was applied after each of the first two weeks of training, and the complete assessment protocol was administered after week 3 (post-test assessment). This took place over three consecutive days: clamp adaptation using scenarios 1 and 2 (day 1); random perturbation and the target-jump task (day 2); and a clamp test under the default map (day 3) to examine if the original operator was preserved.

In the fourth week, we introduced the task-switching manipulation (mixed test) to assess flexibility and context-dependent generalization. Participants completed all three training tasks as before, but the mapping (default or mirrored) was randomly selected on each trial. A clamped feedback probe was administered on days 3 and 6 of this week, with the mapping also randomized with a cycle of 16 trials (see Fig S5a for details). All assessment sessions, and the first training session of weeks 1 and 4, were conducted in the laboratory under the experimenter’s supervision. The remaining training sessions were completed at home.

Two naïve control groups were tested for comparison with the training group in Exp 1. The first control group (n = 17) was included to examine untrained performance on the random-perturbation probe. Targets were drawn from only four locations (45°, 135°, 225°, and 315°) to ensure that participants could produce reasonable baseline behavior with only brief training. To familiarize participants with the mirror mapping, the group completed 40 trials with feedback based on the mirror reversal. This was followed by 20 no-feedback trials and then a 630-trial block with the random perturbations.

The second control group (n = 12) was tested to determine whether a purely visual context cue can, on its own, separate two opposing operators of implicit recalibration. Participants performed movements under the default mapping. We used a target set and trial order matched to the mixed test (see Fig S5b). Importantly, we associated the 15° clamp with a black background and the −15° clamp with a gray background.

#### Experiment 2: Feedforward planning versus feedback control

Exp 2 was designed to determine which aspect or aspects of the training procedure were critical for inducing the reconfiguration of implicit recalibration observed in Exp 1. Participants were randomly assigned to one of two training groups. One group (n = 12) was only trained in the mirrored environment with the center-out reaching task, performing three 120-trial blocks each day (360 total trials/day). The other group (n = 12) was only trained with the continuous tracing task (12 trials per day, with a 5-min break after the first six trials). Time on task was roughly matched across the two groups, approximately 20 minutes per day, and the participants trained for seven consecutive days. Assessments with clamped feedback were administered at pre-test, and after training on days 4 and 7. We only used scenario 1, given that it yields opposite-direction predictions for responses to these clamps under the default and mirror maps.

We also assessed generalization across tasks at the end of session 7. The continuous tracing group was tested on the center-out reaching task, and the center-out reaching group was tested on the continuous tracing task. All participants also completed the target-jump task.

#### Experiment 3: Visuomotor rotation

To examine whether implicit recalibration can also be reconfigured for mappings other than the mirror reversal, we trained a group of participants (n = 12) on a 90° counterclockwise rotation. For a pre-test assessment, participants performed movements to three targets (135°, 225°, and 315°). They were explicitly informed about the nature of the perturbation and instructed to use an explicit aiming strategy, aiming at 45°, 135°, and 225°, respectively, for the three targets. Participants completed a short practice with rotated feedback (5 trials per target), followed by a no-feedback block (15 trials per target). The clamped feedback probe was then administered with the clamp fixed at −45° for all targets. Pilot testing indicated that generating a rotational operator across the whole workspace (0–360°) was highly challenging. We therefore restricted training to a single quadrant, with targets randomly sampled from 115° to 155° in 5° increments.

Participants then completed a two-week training phase with the 90° counterclockwise rotation mapping (6 days per week). Post-test assessments were administered after training on days 6 and 12. Targets were again limited to the 9 locations falling within a wedge spanning 115°–155°, with the −45° clamp applied to the selected target location. To compare the pre-test and post-test assessments, we focused on the 135° target since this lies within the post-test target range. We ended the post-test assessment session with a −45° clamp task under the default map. Participants were informed that the rotational mapping had been removed and were instructed to reach directly to the target. We applied the same targets and clamp as in the assessment under the rotational map.

## Data analysis

All preprocessing and analyses were performed in MATLAB R2024b. For each trial, the cursor position was sampled every frame and converted to polar coordinates (radial distance and angular direction), using the start location as the reference point.

For the center-out reaching task, reach error was defined as the angular difference between the hand angle and the target angle, wrapped to the interval [−180°, 180°]. Trials with a movement time longer than 500 ms or an absolute reach error larger than 45° (wrong-way movements) were excluded, after which residual outliers of each measure (values more than 2.5 SD from the median) were removed. Training performance was summarized on the non-perturbed trials by the mean absolute reach error and the mean reaction time within each session. For the point-to-point reaching task, performance was quantified by the movement duration, defined as the sum of the reaction and movement times averaged across the six reaches of the cycle.

For the continuous tracing task^11^, each 60-s trajectory was reconstructed from the recorded cursor path, resampled onto a uniform grid in time, and transformed with a discrete Fourier transform separately along the horizontal (non-perturbed) and vertical (mirror-reversed) axes. Tracing coherence was quantified, for each axis, as the Pearson correlation between the cursor’s amplitude spectrum and a design vector equal to the target’s stimulus amplitudes at the stimulus frequencies and zero at all other frequencies (horizontal axis: frequencies [7, 16, 34, 49, 67, 91, 121] and amplitudes [100, 100, 100, 50, 50, 30, 30]; vertical axis: frequencies [7, 19, 40, 58, 85, 109, 127] and amplitudes [100, 100, 50, 50, 50, 30, 30], expressed in cycles per 60-s trial). A higher coherence indicates a cursor whose frequency content more closely matches that of the target.

For the clamped feedback assessment, we express adaptation in terms of a mirror-appropriate response (i.e., positive values are changes in hand angle that would be appropriate given the observed error). Hand angles were sign-aligned across the targets of a block. The learning function for each participant was baseline-subtracted using the no-feedback cycles that preceded the clamp phase and corrected for slow drift with a separately measured no-clamp baseline. The magnitude of implicit recalibration was summarized as the mean hand angle over the second half of the clamp phase.

For random-perturbation blocks^33^, implicit recalibration was measured as the trial-to-trial change in hand angle *Δθ* = *θ_n_*_+1_ - *θ_n_*, grouped by the perturbation applied on trial n (±5°, ±10°, ±15°). The strength of adaptive correction was summarized as the difference between *Δθ* at the positive perturbations (+5°, +10°, +15°) and that at the negative perturbations (−5°, −10°, −15°), with positive values again corresponding to corrections in the mirror-appropriate direction.

For the target-jump analysis^16^, trajectories were rotated so the original target lay at 0°. Single-trial trajectories were low-pass filtered (fifth-order Butterworth filter, 20-Hz cutoff, 1-kHz sampling rate), then averaged separately for jump-left and jump-right conditions. The correction function was the signed difference in y-position between these two averages, time-locked to movement onset (right–left for the default map; left–right for the mirror map). Latency was estimated in two steps. First, we identified the earliest time (t*) at which |*Δy*(*t*)| exceeded an absolute threshold of 10 pixels. Second, we backtracked to the earliest prior sample in which the instantaneous slope | *dΔy*/*dt* | first exceeded 0.01 pixels/s in the corrective direction. This sample was taken as the onset of the online error correction.

Improvement across the training phase was assessed using linear mixed-effects models, with log-transformed training day as a fixed effect (capturing the saturating learning curve) and by-participant random intercepts and slopes. One-sample t-tests were used to test whether a measure differed from zero, and paired-sample t-tests to compare measures between the pre-test and post-test assessments. For independent-samples comparisons, Welch’s t-tests were used when the two samples had unequal variances. For the linear mixed-effects models, we reported the fixed-effect slope (β), z value, and p value; for every t-test, we reported the test statistic, its degrees of freedom, the two-tailed p value, the partial η² effect size, and the Bayes factor in favor of the alternative hypothesis (BF₁₀). The significance level was set at two-tailed p < 0.05.

## Author contributions

T.W., J.A.T., and R.B.I. contributed to the conceptual development of this project. T.W. and Y.L. implemented the experiment and collected the data. T.W. and Y.L. analyzed the data and prepared the figures. T.W. and Y.L. wrote the initial draft of the paper. T.W., J.A.T., and R.B.I. were involved in the editing process.

## Acknowledgments

R.B.I. was funded by grants NS116883 and NS105839 from the National Institutes of Health (NIH). J.A.T. was funded by grants NS131552 and NS134754 from the NIH.

## Supplementary Material

### Supplementary Methods

The twelve Exp 1 participants were tested in three groups, collected at different times as the paradigm was refined, and the groups followed somewhat different training schedules. The principal difference was the duration of training. One group (n = 3) trained under the mirror reversal for four weeks (20 daily sessions). The other two groups each trained for three weeks (15 daily sessions) before completing the additional task-switching week (the mixed test). The groups also differed in the clamp used to probe implicit recalibration: one group (n = 4) used a ±15° clamp, whereas the other two groups (n = 8) used a ±30° clamp. All other aspects of the training and assessment protocol were common to the three groups. Because every measure was referenced to each participant’s own pre-training baseline, these scheduling differences do not affect the within-participant comparisons reported in the main text.

### Supplementary Results 1: Implicit recalibration generates trial-by-trial adaptation under mirror reversal

We used clamped feedback in our initial demonstration that implicit recalibration can adapt to a novel sensorimotor map following extended training. As a second test, we examined a subset of the Exp 1 participants (n = 9) with a set of different perturbations that randomly varied in size and direction from trial to trial. Although the participants had control over cursor position, they were informed that the perturbations were random and instructed not to re-aim since “correcting” for these errors would not improve performance. As shown in previous studies, when perturbations vary randomly from trial to trial, participants do not explicitly re-aim under such conditions whereas implicit recalibration continues to respond automatically from trial to trial^33,55^. Importantly, the predicted change in hand angle is quite different depending on whether this system operates on the default map or the mirror map. If implicit recalibration remains anchored to the default map, the SPE, the discrepancy between the hand and cursor positions, would be relatively constant (Fig S2a, left). In principle, this would drive a gradual, accumulated drift away from the target^37^ (see Fig 2c). By randomizing the direction of the perturbation, we focused on the trial-by-trial corrections, predicting that they would be negligible across perturbation sizes if the implicit recalibration system operates on the default map. In contrast, if the implicit system operates under the mirror map, the SPE will reflect the target error, since the sensory prediction now aligns with the target. As such, the corrections will be correlated in sign and magnitude with the perturbation.

The results clearly showed that after extended training, implicit recalibration was responsive to the perturbations, suggesting that it had now become reconfigured to the mirrored sensorimotor map (Fig S2c, t(8) = 8.1, p < 0.001, ηp² = 0.89, BF₁₀ = 597). The corrections were sign-dependent, shifting in the same direction as the perturbation, a response that would appropriately compensate for the error. Moreover, the magnitude of the response scaled with the size of the perturbation. To confirm that these findings were the result of the prolonged training, we tested a separate naïve control group (n = 17) with the same random perturbations. The single-trial corrections for this control group showed no dependency on the direction or size of the perturbation (t(16) = 1.6, p = 0.14, ηp² = 0.13, BF₁₀ = 0.70), indicating that for these participants, implicit recalibration was based on the default map. A direct comparison of the two groups showed that the magnitude of the average correction was much larger in the trained group than in the untrained group (Fig S2d, Welch’s t(14.2) = 6.0, p < 0.001, ηp² = 0.72, BF₁₀ = 3.4×10³). The finding that RT was uncorrelated with perturbation size (Fig S2b) is consistent with the hypothesis that the perturbation-scaled correction in the trained group was not due to an explicit re-aiming strategy^36^.

### Supplementary Results 2: Online error correction becomes automatized through extended training

Beyond generating the correct feedforward command, another hallmark of skilled performance is the ability to rapidly make online adjustments to an ongoing movement. We assessed this ability before and after training with a task in which the target unexpectedly changes position mid-movement (Fig S3a). Approximately 100 ms after movement onset, the target jumped 30° clockwise or counterclockwise, or remained stationary. We administered this probe when the feedback was mirrored in the pre-test and post-test phases of Exp 1, as well as in a baseline condition in which the feedback was veridical (default map).

Under the default map, corrective responses were initiated rapidly, with a single lateral shift in the trajectory that started around 200 ms after the target jumped (Fig S3b–c). In the pre-test under the mirror reversal, participants showed a biphasic deviation in the trajectory^16^. The first response was in the opposite direction of the jump, a maladaptive response for a mirror environment, with a similar onset compared to the response under a default mapping. This early response was then overwritten by a correction in the opposite direction, steering the cursor back toward the target. This replicates the pattern described previously.

After extended training in the mirrored environment, this biphasic response collapsed into a single, rapid correction in the appropriate direction. This correction began much earlier than the adaptive, secondary response observed prior to training (t(11) = 3.6, p = 0.004, ηp² = 0.54, BF₁₀ = 12), yet was still slower than the correction to a target jump observed at baseline under the default map (t(11) = 4.9, p < 0.001, ηp² = 0.69, BF₁₀ = 80). This intermediate latency suggests that the proper correction is initially slow and cognitively mediated but becomes progressively automatized with practice. Together with the implicit-recalibration results, extended mirror training appears to reconfigure both the implicit recalibration system and the online-correction process.

### Supplementary Results 3: Generalization of training on the center-out and tracing tasks

To understand the training consequences of continuous tracing and center-out reaching, we compared the two training groups in Exp 2 on three different tasks after training, comparing their performance to that observed in Exp 1 on the first day of training in the mirrored environment.

On the center-out reaching task, both groups improved substantially relative to the comparison group (Fig S4a), an unsurprising result given that continuous tracing also requires feedforward planning. As expected, the center-out training group performed better at post-test, as evident in throughput, a composite measure accounting for the speed–accuracy trade-off (t(21.2) = 3.0, p = 0.006, ηp² = 0.31, BF₁₀ = 7.7). For the continuous tracing task, the center-out group was markedly poorer on both axes (Y: t(21.8) = 4.3, p < 0.001, ηp² = 0.47, BF₁₀ = 95; X: t(17.7) = 3.0, p = 0.008, ηp² = 0.33, BF₁₀ = 6.8), indicating that tracing imposes demands not captured by center-out reaching.

The critical comparison is the target-jump task, which is performed in a center-out configuration but requires rapid feedback-based correction. We note that the two groups showed similar levels of improvement in terms of correction latency (Fig S4c, d; t(21.8) = 0.21, p = 0.84, ηp² = 0.0020, BF₁₀ = 0.38), with both correcting ∼105 ms earlier than the slow response in the pre-test (center-out: t(17.6) = 3.5, p = 0.003, ηp² = 0.40, BF₁₀ = 13; tracing: t(17.9) = 3.1, p = 0.007, ηp² = 0.34, BF₁₀ = 6.9). Thus, on this measure, center-out training enhanced feedback-based control and learning acquired through tracing transfers to the discrete center-out configuration. Critically, despite this equivalent online-correction ability, the two groups showed distinct patterns of implicit recalibration at post-test. This dissociation indicates that mirror-appropriate online correction is neither required for, nor the driver of, reconfiguration of implicit recalibration.

---

**Figure S1.**
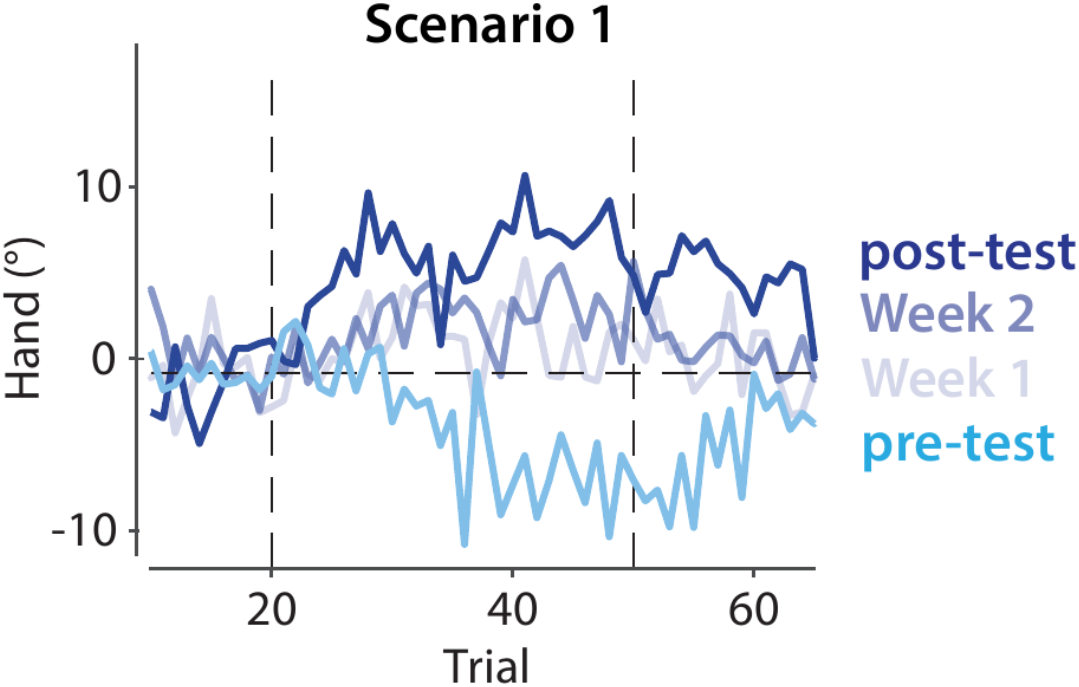
Implicit recalibration is gradually reconfigured during mirror training in Experiment 1. We used the scenario 1 perturbation to probe implicit recalibration after each week. After one week of training, the inappropriate response based on the default map was suppressed. The appropriate response based on the mirrored map became more pronounced over subsequent weeks.

**Figure S2.**
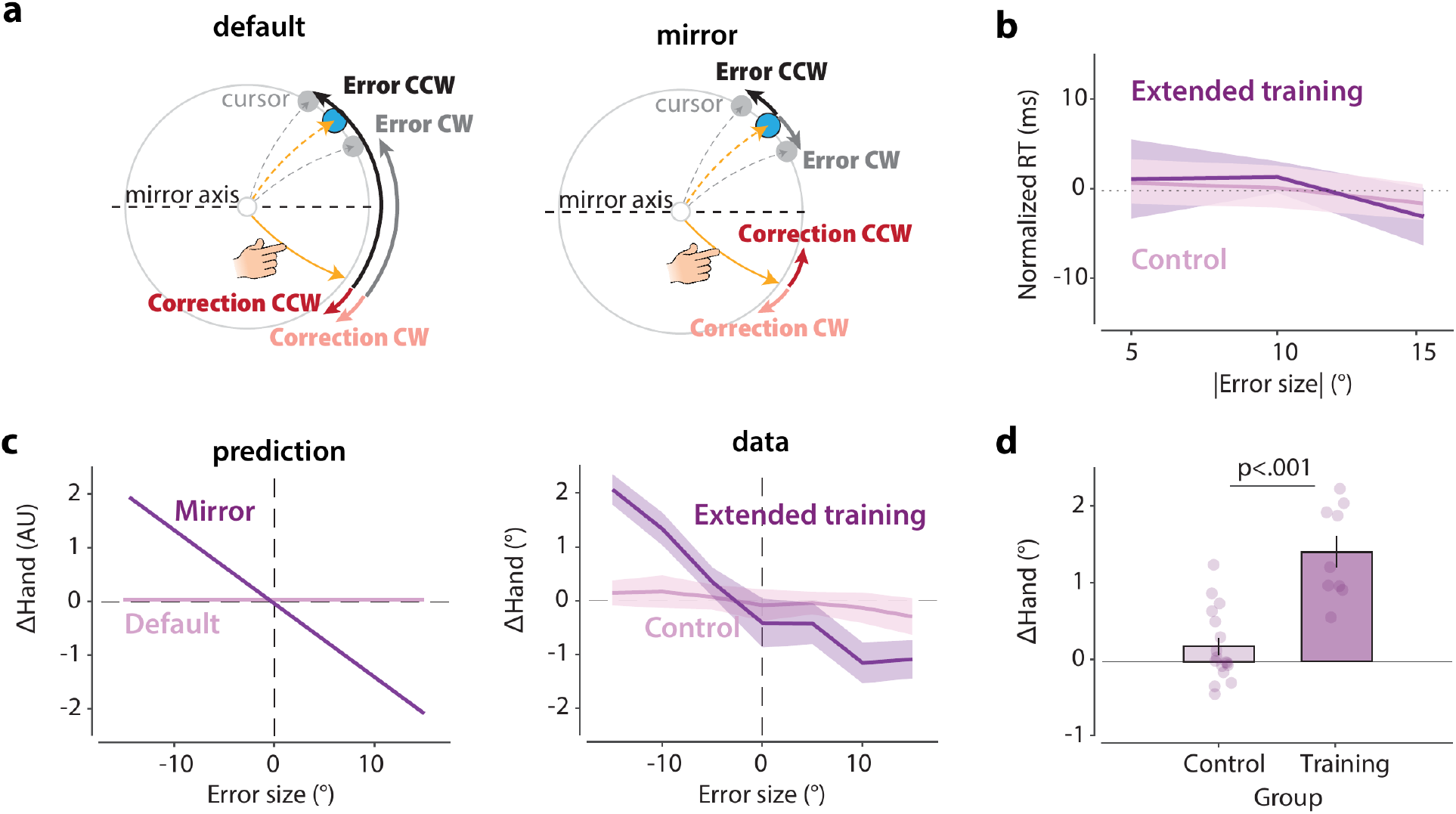
Implicit recalibration corrects errors in a trial-by-trial manner after prolonged training. (a) Probe operation of implicit recalibration in response to hand-contingent perturbations that vary in size and direction. The target position was held constant for seven consecutive trials before switching to a new location. For a given target, each of the six nonzero perturbations (±5°, ±10°, or ±15°) was applied once in random order, followed by an unperturbed (0°) trial. The dashed yellow line indicates the unperturbed cursor trajectory (assuming reach has zero error) and dashed gray lines depict two perturbations (±15°). Left: if the implicit system operates under the default map, the two perturbations would be perceived as similar error signals, resulting in similar corrections. Right: if the implicit system operates under the mirror map, the two perturbations generate error signals with opposite signs and would result in corrections of opposite directions. (b) Normalized reaction time as a function of perturbation size on the previous trial. RT did not increase with perturbation size. (c) Single-trial error correction as a function of the size of the random perturbation. Left: Predicted behavior under the default and mirror maps. For the former, the perceived error is relatively constant. For the mirror map, error correction should be sign-dependent and scale with error size. Right: Error correction functions in the extended-training group (Exp 1) and the naïve control group. The results match the predictions of the mirror and default mappings, respectively. (d) Mean error correction per trial. Positive values indicate corrections aligned with the mirror-reversal mapping.

**Figure S3.**
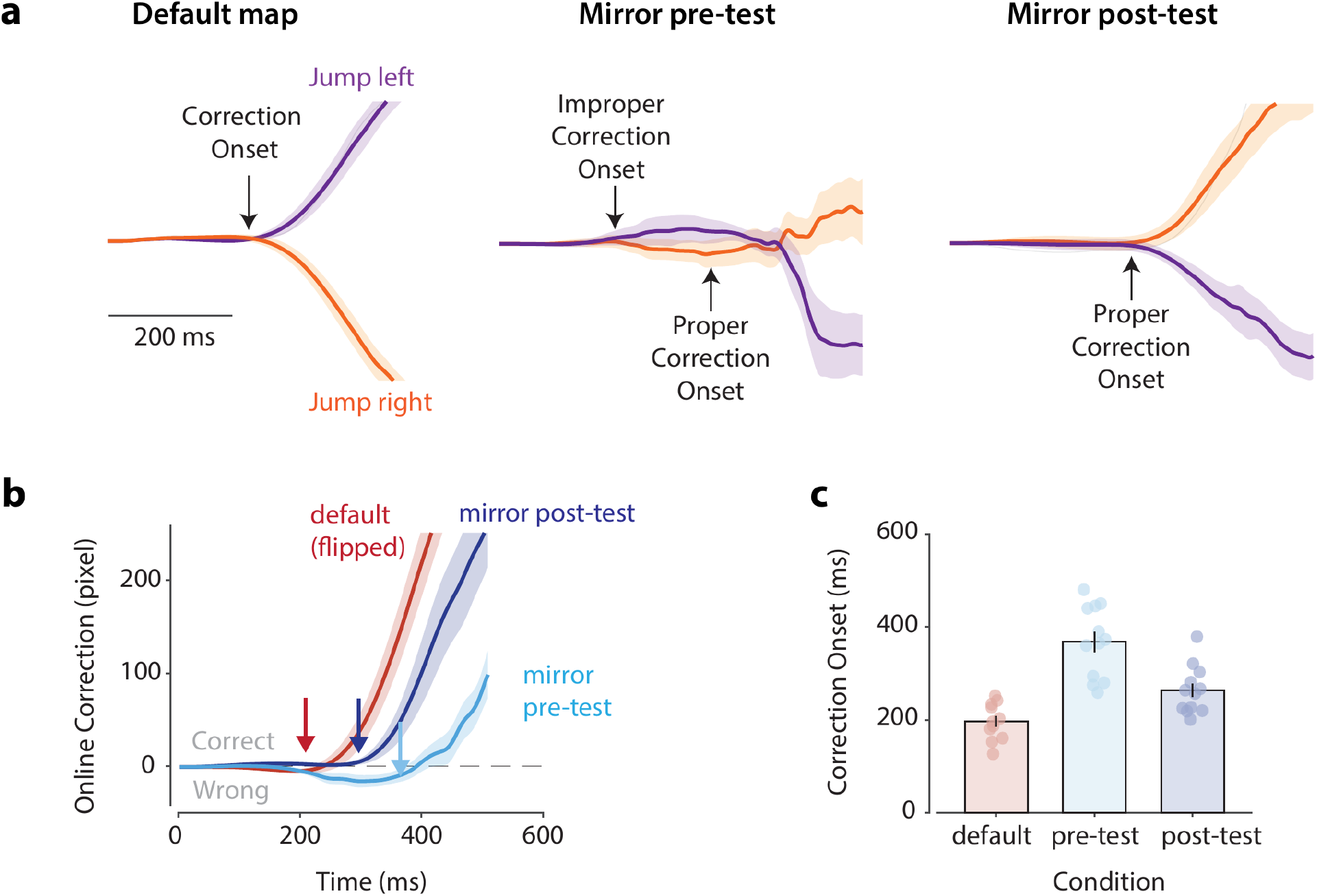
Online correction speeds up after mirror training. (a) Movement trajectories following a target jump under the default and mirror mappings. Trajectories are grouped by jump direction, with the original target aligned to the x-axis. (b) Time course of online correction, computed as the difference in y-position between jump-left and jump-right trajectories. Positive values indicate a correction in the appropriate direction. In the mirror-reversal pre-test, there was a biphasic pattern composed of an inappropriate initial response followed by a correction in the proper direction. In the default mapping and the mirror post-test, only a single correction was observed, one that brought the cursor back toward the target. Arrows mark the detectable correction onset for each condition. (c) Correction onsets by condition. The latency of this response (mirror-reversed mapping) remained slower than that observed at baseline (default mapping) in both pre-and post-test assessments.

**Figure S4.**
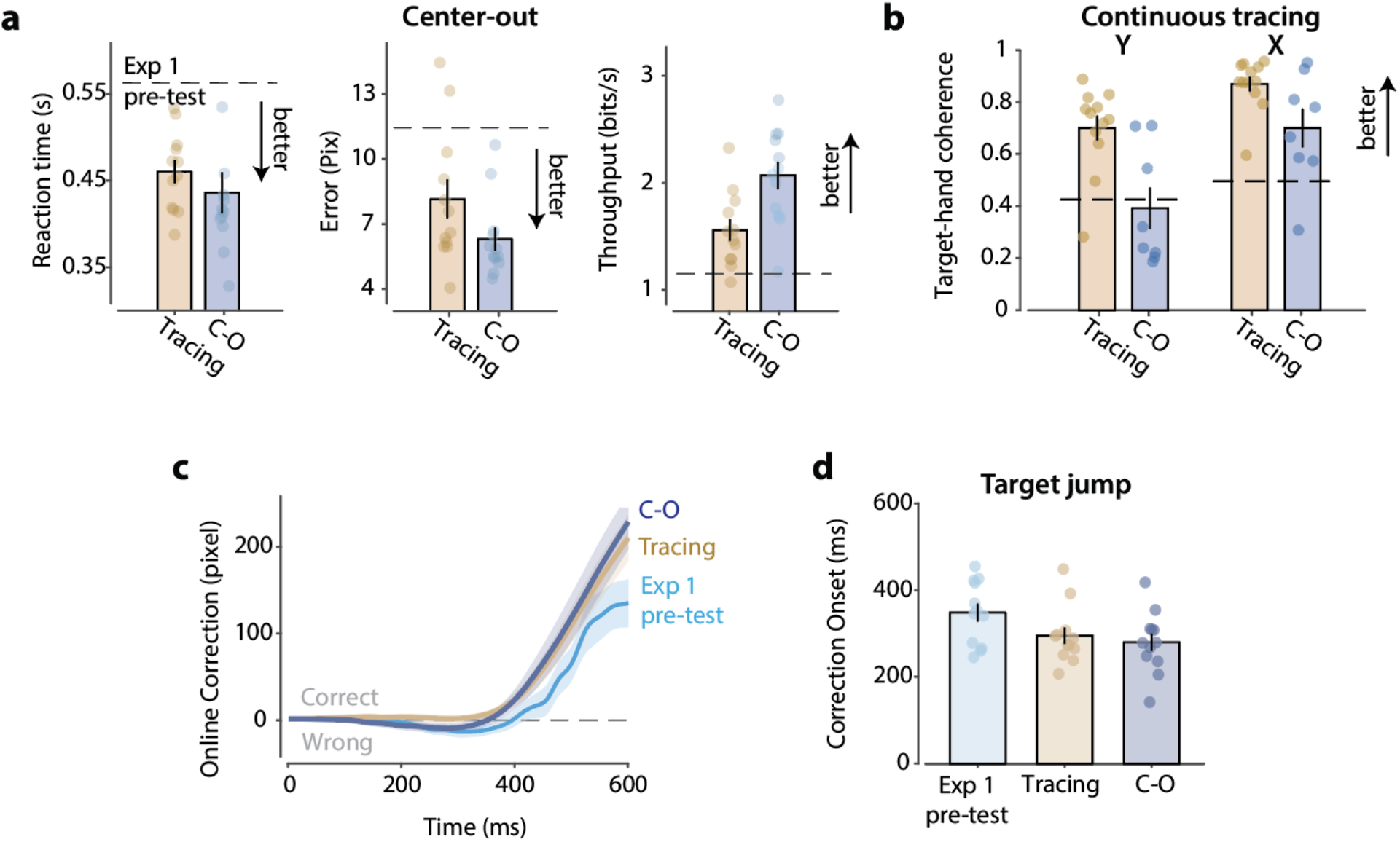
Generalization of learning across tasks in Experiment 2. (a) Performance on a center-out reaching task for the tracing and center-out training groups on measures of reaction time, absolute error, and throughput (a composite measure accounting for the speed–accuracy trade-off). (b) Performance on a continuous tracing task, quantified as target–hand coherence (see Methods) along the x-axis (non-perturbed) and y-axis (mirror-reversed). (c) Lateral deviation of the hand in response to a target jump. (d) Onset latency of this deviation for the three groups.

**Figure S5.**
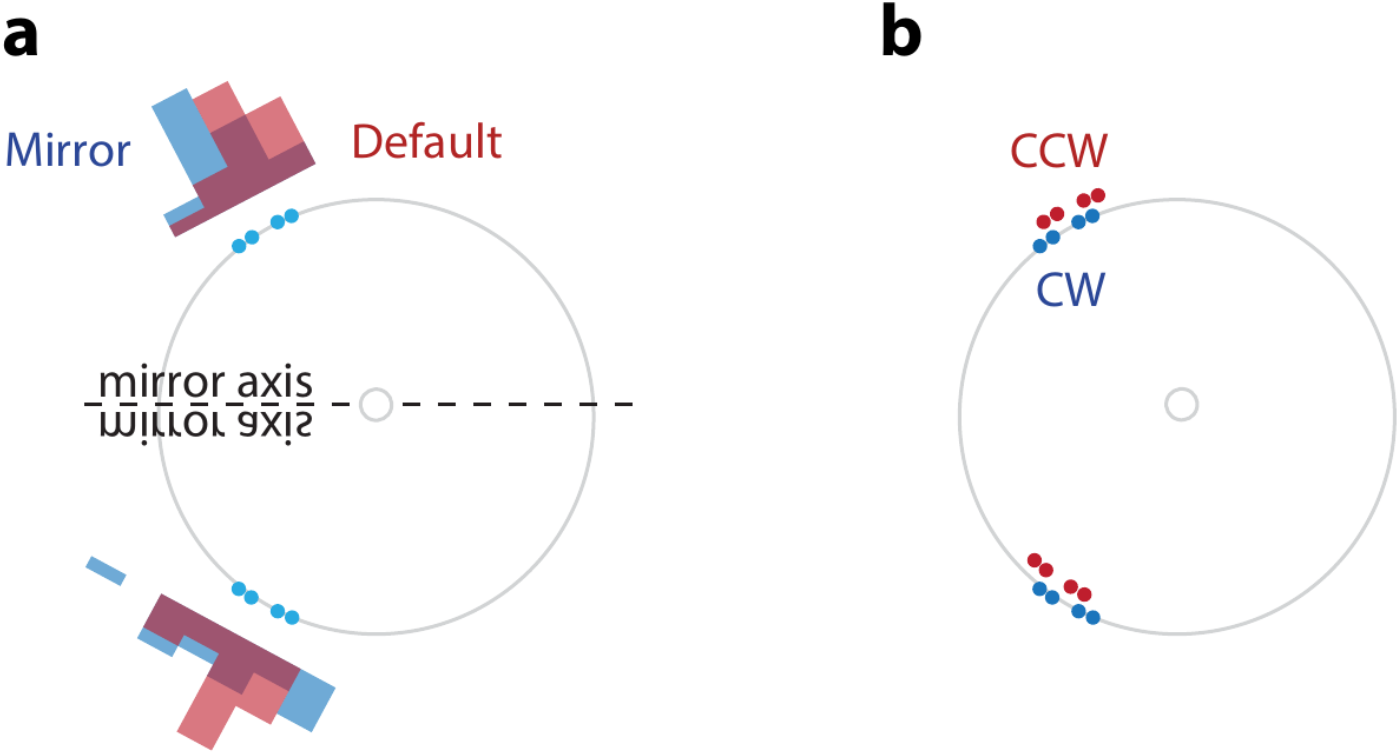
Target configuration for the mixed test. (a) To ensure that reaching directions largely overlapped between the mirror and default maps, we used two sets of targets that were symmetric across the mirror axis. Because the control policy may not be perfect and participants’ movement directions may be slightly off from the mirrored target in the mirror condition, we jittered the targets within a 16° range. The histogram shows that the distributions largely overlap for baseline reaching directions for the mirror (blue) and default (red) mappings. (b) As a control task, we trained participants on a dual-adaptation paradigm using CW and CCW rotational perturbations. The CCW targets were identical to those used in panel a. For the CW condition, we rotated all targets by 5° to introduce a small mismatch between the mean reaching/aiming directions for CW and CCW conditions. This was done to compensate for the mismatch between the reaching distributions for the default and mirrored mapping conditions shown in panel a.

